# Three-dimensional body reconstruction enables quantification of liquid consumption in small invertebrates

**DOI:** 10.1101/2024.06.14.599002

**Authors:** Henrique Galante, Tomer J. Czaczkes, Massimo De Agrò

## Abstract

Quantifying feeding patterns provides valuable insights into animal behaviour. However, small invertebrates often consume incredibly small amounts of food. This renders traditional methods, such as weighing individuals before and after food acquisition, either inaccurate or prohibitively expensive. Here, we present a non-invasive method to quantify food consumption of small invertebrates whose body expands during feeding. Using the markerless pose estimation software DeepLabCut, we three-dimensionally track the body of Argentine ants, *Linepithema humile*. Using these extracted markers, we developed an algorithm which computationally reconstructs the ant’s body, directly measuring volumetric change over time. Moreover, we provide measures of accuracy and quantify the ant’s feeding response to a range of sucrose concentrations, as well as a gradient of caffeine-laced sucrose solutions. Small invertebrates are often prolific invasive species and disease vectors, causing significant ecological and economical damage. Understanding their feeding behaviour could be an important step towards effective control strategies.

## Introduction

A fundamental aspect of science is measuring quantities. However, all devices are subject to error, making it impossible to obtain truly exact measurements. This becomes increasingly difficult as scale decreases (Jenkins & Cook, 2004; Deshpande *et al*., 2014). In biology, small animals such as mosquitoes, flies or ants often ingest volumes in the nanolitre range (Paul & Roces, 2003; Wong *et al*., 2008; Jové *et al*., 2020). This is problematic when studying animal behaviour as equipment capable of measuring such small volumes with both precision and accuracy is prohibitively expensive (Councill *et al*., 2021). Additionally, such devices often work exclusively under tailored conditions which are seldomly possible when studying natural behaviours (see Calisi & Bentley, 2009 for a review). Yet, feeding behaviour provides valuable insights into an animal’s perception of the world, elucidating their preferences and cognitive capabilities (Kim & Smith, 2000; Oberhauser, Koch & Czaczkes, 2018). Additionally, feeding is often linked to nutritional status and can be a measure of animal welfare (Vaudo *et al*., 2016; Carvajal-Lago *et al*., 2021). Finally, feeding is also particularly important in pharmacological interventions (Devineni & Heberlein, 2009; Vinauger *et al*., 2018) and can play a crucial role in the development of effective invasive species control methodology (Nigg *et al*., 2004; Carrasco *et al*., 2019; Galante & Czaczkes, 2024; Galante *et al*., 2024). Thus, an affordable and reliable method for quantifying feeding behaviour in small invertebrates would be extremely valuable.

Currently, one of the most common methods of measuring feeding in small invertebrates is the capillary feeder assay. This involves quantifying the depletion of liquid food from a glass capillary, thereby providing a measure of the ingested volume (Ja *et al*., 2007). However, this approach requires the animal to drink upside-down from a vertical capillary, potentially restricting the method’s applicability to animals who can do so. Moreover, the food used is often dyed to facilitate the measurement of depletion from the capillary, which can impact feeding behaviour by altering food consumption (Shell *et al*., 2021). Lastly, since the assay directly measures food depletion rather than the actual volume ingested by the organism, it is highly sensitive to evaporation and spillage (Diegelmann *et al*., 2017).

Gravimetric approaches are a widely used alternative, where an individual, or the food source, is weighed before and after feeding to quantify consumption (Rotheray, Osborne & Goulson, 2017; Straw & Brown, 2021). However, for small animals, this requires an extremely sensitive scale, which is both expensive and cumbersome to work with. Furthermore, measuring the weight of a dynamic object is complex, often requiring individuals to be anaesthetised before and after the feeding event, which may alter their behaviour and could influence food consumption (Mailleux, Deneubourg & Detrain, 2000; Poissonnier, Jackson & Tanner, 2015; Gooley & Gooley, 2023). More importantly, weighing does not provide information about consumption rate patterns during feeding and is strongly influenced by external factors, such as liquids which were not ingested adhering to the individual’s body.

Another popular alternative takes advantage of dyes, whether food-grade or fluorescent, or of other chemical tracers (Nigg *et al*., 2004; Řehoř *et al*., 2014; Wu *et al*., 2020; Sakuma & Kanuka, 2021). While these methods may offer easy quantification, they often come with steep equipment costs and provide indirect measures of food ingested through spectral analysis or similar processes. These chemicals can potentially alter feeding behaviour, and often interact with other food components, such as protein, which can interfere with quantification (Marmé *et al*., 2003; Baltiansky *et al*., 2021). Additionally, dyes can stain the individual’s tissue and methodology often requires the sacrificing of the individual post-consumption.

Dedicated methods have previously been developed to address the complexity of quantifying feeding in small invertebrates. The flyPAD system, for example, can measure capacitance changes when an individual touches the food source (Itskov *et al*., 2014; Henriques-Santos, Xiong & Pietrantonio, 2023). This method has proven useful for characterising feeding behaviour, offering an indirect measure of consumed solution based on the time spent touching the reward, assuming linear consumption rates. However, this system is costly and requires rigorous maintenance and cleaning. Moreover, the non-open-source nature of the original hardware and software presents a limitation. Importantly, it also requires individuals to be anesthetised and placed in a small artificial chamber, which could alter their behaviour and food consumption (Mailleux, Deneubourg & Detrain, 2000; Poissonnier, Jackson & Tanner, 2015; Gooley & Gooley, 2023).

Here, we aim to address the limitations of currently available methods, developing a high-resolution, high-throughput and non-invasive system to quantify food consumption of small invertebrates. The body of many insects (Hymenoptera, Diptera, Hemiptera, Lepidoptera), some arachnids (Ixodida, Araneae, Opiliones) and worms (Annelida, Nematoda) visibly expands as they ingest food. Quantifying this expansion over time allows for measures of volumetric increase, and consequentially of liquid ingested. A similar approach has previously been developed by approximating volume consumption from two-dimensional tracking of body shape (Sola & Josens, 2016; Hol, Lambrechts & Prakash, 2020). We expand on this, using the well-documented open-source software DeepLabCut (Mathis *et al*., 2018; Nath *et al*., 2019), by tracking body shape in a three dimensional space. Using these extracted markers, we developed an algorithm which computationally reconstructs the animal’s body over time, thus directly measuring volumetric changes. In this way, we eliminate the error associated with approximating volume from area, which occurs when using a two-dimensional system for volume estimation (Sola & Josens, 2016; Hol, Lambrechts & Prakash, 2020). This method allows animals to move freely, without the need for anaesthesia or the use of dyes. Furthermore, it not only directly quantifies the volume of food ingested but also provides consumption rates over time. Understanding feeding patterns provides essential insights of animal behaviour and can be crucial for the study of disease vectors and transmission as well as the development of invasive pest control methodology.

## Materials and Methods

The system we developed takes advantage of the fact that some animals’ bodies expand during feeding to accommodate for the ingested food, enabling us to quantify feeding patterns. The process can be divided into five steps, detailed below. First, the user should build a setup, adapted to the animal being studied, which is capable of recording the feeding events with sufficient quality (Step 1). Next, using DeepLabCut three-dimensional pose estimation software (Mathis *et al*., 2018; Nath *et al*., 2019) key points of the individual’s body are extracted from the videos at each time point (Step 2). Once these three-dimensional coordinates are obtained, volumetric changes of the tracked body over time can be quantified using one of the seven methods proposed (Step 3). Finally, our simple graphical user interface (GUI) allows the user to select the initial and final times of the feeding event (Step 4) and fit a model to the individual’s volume over time (Step 5). This approach provides both the total amount of food ingested during the feeding event and the individual’s consumption rate.

### System specifications

#### Step 1 - Physical setup

In most cases, a basic recording configuration, consisting of two perpendicular cameras, capturing a top-view and side-view of the features of interest, should suffice. Nevertheless, additional cameras, placed at varying angles may be required, particularly if recording subjects mid-flight (Maya *et al*., 2023; Håkansson *et al*., 2024), for example. Regardless of the number of cameras used, it is imperative these are stereo-configured and time-synchronized to ensure overlapping views and simultaneous visibility of features of interest across multiple cameras, allowing for the feature to be reconstructed in a three-dimensional space. Once the system is built, it is important to take calibration images, for example by using a checkerboard. The calibration step will track known points across all cameras simultaneously, computing intrinsic (optical centre and focal length of the camera) and extrinsic (camera location in the three-dimensional space) parameters for each camera. This step is critical for successful triangulation, as it combines different views of the same point, in order to determine its location in the three-dimensional scene. For further details we recommend consulting the DeepLabCut user guide for 3D pose estimation (Mathis *et al*., 2018; Nath *et al*., 2019). The specific physical setup we developed to validate the volume estimation method with ants is described below.

#### Step 2 - Three-dimensional pose estimation

DeepLabCut version 2.3.8 (Mathis *et al*., 2018; Nath *et al*., 2019) was used to track points of interest in 3D. Two distinct DeepLabCut networks, one for each camera, were trained and later used during experimental validation and application. Each network was trained on 15 manually labelled frames from 28 videos (840 labelled frames and 14 280 points labelled in total). The two networks tracked 17 points each, 15 points following anatomical landmarks (Figure 1) in the animal’s body (in this case, ants), and two reference points on the feeding platform (described below) allowing for the standardisation between pixels and millimetres (1 px = 1.38 ± 0.11 mm, N = 538). The network trained on recordings from camera A was trained to 100 000 iterations, with a train error of 3.59 pixels and a test error of 5.73 pixels. The network trained on recordings from camera B was trained to 110 000 iterations, with a train error of 3.75 pixels and a test error of 3.74 pixels. All videos were recorded at 30 frames per second and cameras were stereo calibrated using a 10x8 checkerboard. The output of each video triangulation consisted of a file containing the three-dimensional cartesian coordinates of each of the 17 points tracked at each frame of the video, alongside a likelihood value indicating the model’s confidence in the point coordinate estimation. Points with a likelihood below 60% were discarded during triangulation.

**Figure 1.**
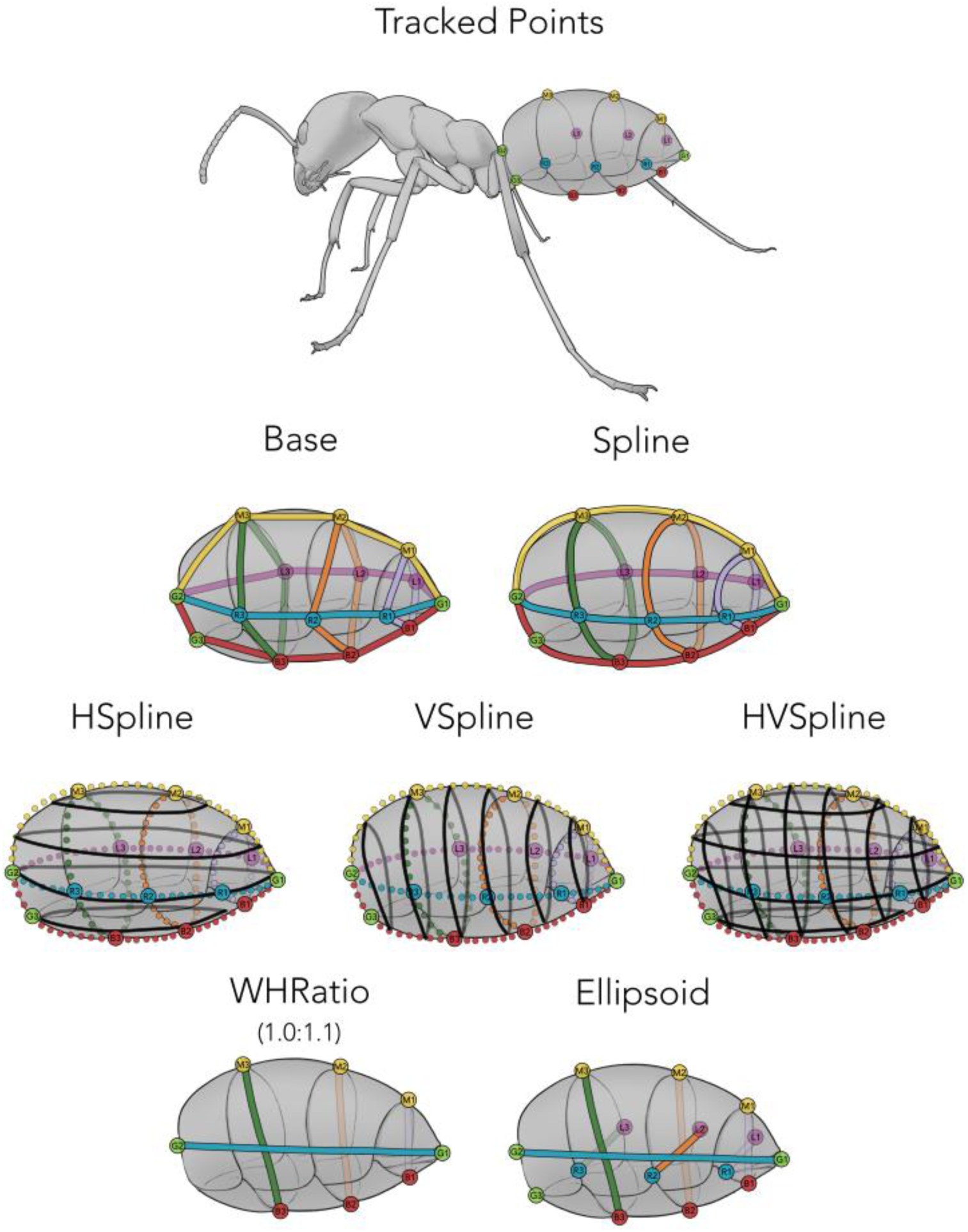
Anatomical landmarks tracked using three-dimensional pose estimation and how these were used to estimate volume using seven approaches. The **Base** method simply connects the 15 tracked points using straight lines. **Spline** connects these points by fitting 100 interpolated points to curves. **HSpline** and **VSpline** further interpolate points along horizontal and vertical connections of the previously generated points, respectively. **HVSpline** combines both horizontal and vertical connections. **WHRatio** uses a species-specific width to height ratio to approximate the skeleton to an ellipsoid, and **Ellipsoid** does so but without using a predefined ratio.

#### Step 3 - Volume estimation methods

Python version 3.7.13 (Rossum & Drake, 2009) was used to process the triangulated points obtained from DeepLabCut. The algorithm developed loads the cartesian coordinates for each recording, uses the reference points to scale values from pixels to millimetres, and estimates the volume of the target feature using seven different approaches (Figure 1).

The first approach, which we named the **Base** method, quantified the volume of the convex hull: the simplest polygon encompassing a set of points, in this case formed by the 15 tracked points around the ant’s gaster. While this method is simple and requires no assumptions of the shape of the animal’s body, it creates straight lines between the points rather than curves. Thus, in order for it to represent the true volume of the object, one would need to track an infinite number of points describing its shape. Yet, labelling points is time consuming and pose estimation is computationally costly. Thus, the **Spline** approach expands on this simple method, by interpolating 100 points for each of the seven splines (smooth, bendable lines used to connect points in a continuous, curvilinear manner) created by the object’s main axes (highlighted in different colours in Figure 1), thus creating a more detailed skeleton, at the cost of assuming the object is curvilinear.

Increasing the level of complexity, the **HSpline** and **VSpline** methods further interpolate points along horizontal and vertical connections of the previously generated points, respectively, creating an even fuller skeleton without the need for tracking more points. The **HVSpline** method simply combines both of these methods by filling the skeleton both vertically and horizontally.

For validation purposes, we include the **WHRatio** method, as described by Sola & Josens (2016). This method approximates the shape of an ant’s gaster to an ellipsoid, using a two-dimensional image and a species-specific width-to-height ratio.

Finally, the **Ellipsoid** approach approximates gaster shape to an ellipsoid, but without using a predefined ratio. Instead, it takes the maximum width and the maximum height values from the tracked points.

#### Step 4 and 5 - Measuring ingested volume and consumption rate

To allow the user to verify if the tracking and volume estimation worked as expected, we designed a simple graphical user interface (GUI). When launched, this GUI plots a graph of the estimated volume at each time point with a slider which can be adjusted to a frame of interest. By moving the slider, the user can select a frame of the video. This is then shown for both cameras and a 3D plot of the object’s reconstruction (in our case, the ant gaster) is also displayed for that frame (Figure 2). Additionally, since the recordings do not necessarily begin and end with the feeding event, we manually select two time points which represent its beginning and end time. To facilitate this, the GUI allows the user to visualise the data being analysed and identify the time at which the individual started and stopped feeding. The indexes of these frames are then automatically saved for later use and the user can choose to inspect a new video or resume the process at a later time. The GUI automatically identifies which videos have not yet been inspected and appends the new data to the same file, allowing the user to seamlessly distribute the process across multiple sessions. In order to quantify feeding behaviour, we extract two main variables from each recording: crop load, a measure of how much food was ingested, and consumption rate, a measure of how fast this food was consumed over time. Using the manually selected start and end points of the feeding event and reducing noise by applying a rolling average filter for each five seconds, we fit a linear regression to the estimated volumes over time. The slope of the linear regression is taken as the consumption rate of that individual, and the regression equation used to calculate the initial and final volume from which crop load is then calculated.

**Figure 2.**
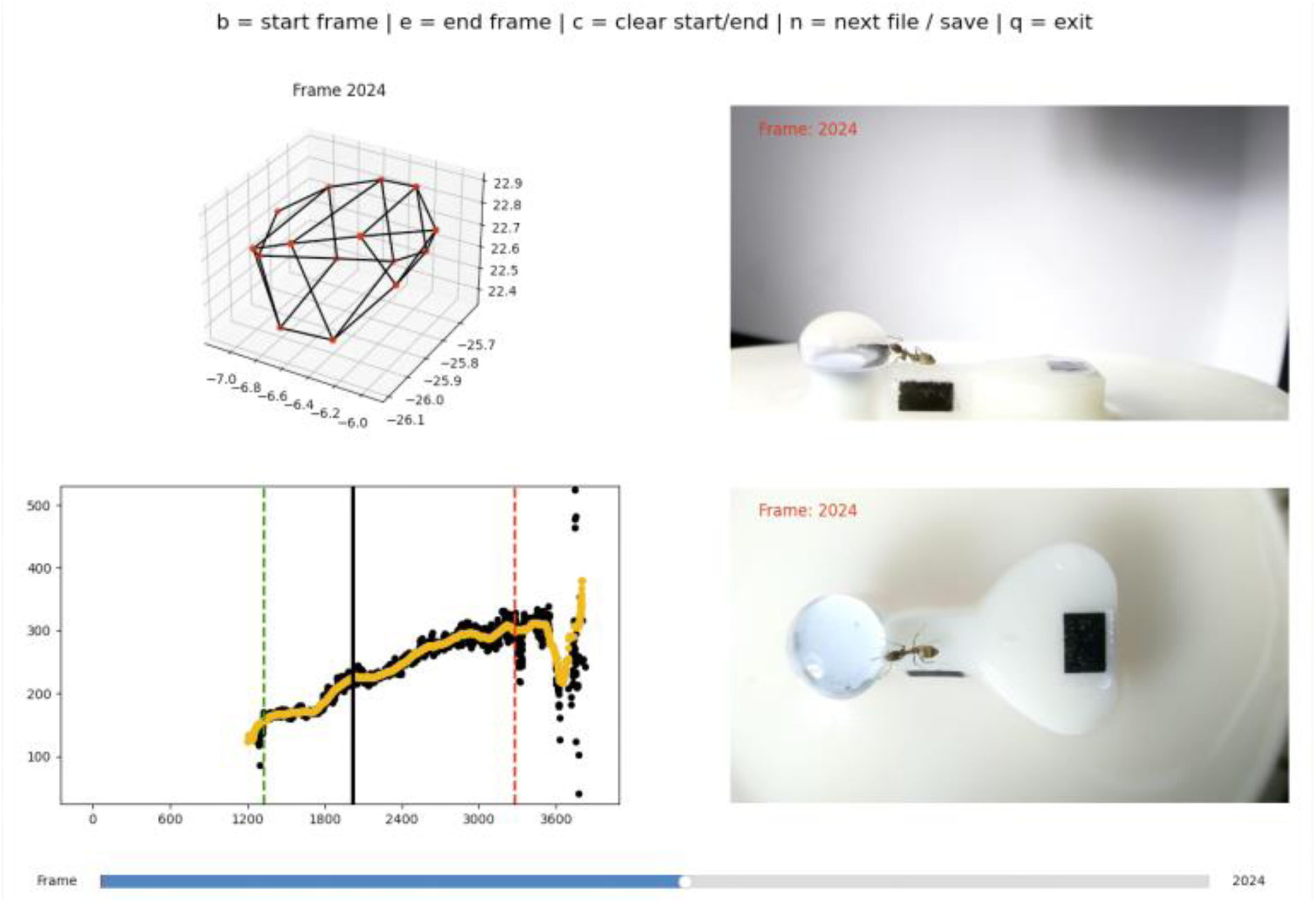
Graphical user interface (GUI) designed to facilitate manual selection of the start (green dashed line) and end point (red dashed line) of the feeding event. The GUI displays the volume over time interactive plot, with points in black showing the raw volume estimates and those in orange a rolling average of these values. By moving the slider, the user can select a frame of interest, displaying it in each camera as well as a 3D plot of the tracked points. This allows the user to validate both the three-dimensionally tracked points and the volume calculations. The manually selected frames are automatically saved and the GUI allows the user to analyse recordings in multiple sessions, resuming from the last recording analysed.

### Quantifying feeding patterns of Argentine ants

In order to validate and apply the developed algorithm, we conducted three experiments using Argentine ants, a globally invasive species. Firstly, we assessed the method’s accuracy by measuring its agreement with gravimetric estimates. To do this, starved ants were initially weighed, then recorded whilst ingesting food, and once full, weighed again. Secondly, we fed ants a gradient of sucrose solutions. As sucrose concentration increases, so does the solution’s viscosity. Ants are expected to prefer and consequently ingest more food when the sucrose concentration - and thus the energy provided by the solution - is higher. However, the high viscosity of these solutions is expected to decrease the rate at which the ants can ingest the food (Sola & Josens, 2016; Fujioka, Marchand & LeBoeuf, 2023). Lastly, we fed the ants a range of caffeine-laced sucrose solutions in order to assess their preference or aversion for them. Caffeine has been reported to decrease foraging times, likely due to its beneficial cognitive effects (Galante *et al*., 2024). Nevertheless, understanding if ants will be attracted or repelled to baits containing caffeine, consuming them at different rates and/or absolute amounts, is crucial if the addition of additives such as caffeine becomes common practice in invasive ant management.

#### Colony maintenance

*Linepithema humile* (Mayr, 1868) were collected from Portugal (Proença-a-Nova) and Spain (Girona) between April 2021 and April 2022. Ants were split into colony fragments (henceforth colonies), containing three or more queens and 200-1000 workers, kept in non-airtight plastic boxes (32.5 x 22.2 x 11.4 cm) with a plaster of Paris floor and PTFE coated walls. 15mL red transparent plastic tubes, partly filled with water, plugged with cotton, were provided as nests. Ants were maintained on a 12:12 light:dark cycle at room temperature (21-26 °C) with *ad libitum* access to water. Between experiments, ants were fed *ad libitum* 0.5M sucrose solution and *Drosophila melanogaster* twice a week. During experiments, ants were fed once a week and deprived of carbohydrates for four to five days prior to testing, ensuring high foraging motivation. Experiments were conducted between January and October 2022 using 571 ants from nine colonies.

#### Solutions

Sucrose solutions (Südzucker AG, Mannheim, Germany) following a geometric sequence ranging from 0.125M to 2M were used as treatments. Caffeine (CAS 58-08-2) was obtained from Sigma-Aldrich (Taufkirchen, Germany). 1M sucrose solutions mixed with different caffeine concentrations were used as treatments. Caffeine concentrations were chosen based on previous reports of their effects on Hymenopterans. Caffeine solutions ranging from naturally occurring concentrations (Singaravelan *et al*., 2005; Couvillon *et al*., 2015) to those reported as the LD_50_ of honeybees (Detzel & Wink, 1993) were used: 25ppm (0.13μmol ml^-1^), 250ppm (1.29μmol ml^-1^) and 2000ppm (10.30μmol ml^-1^), respectively.

#### Experimental validation – Accuracy measurements

Method accuracy was measured by both weighing ants before and after drinking (N = 104), and by feeding them a 138nL drop of 0.125M sucrose solution created by a nanolitre microinjection system (N = 10) whilst simultaneously recording them using our validation setup (described below). The difference between these and the volume estimation for each individual were then used as a measure of accuracy. All weight measurements were performed using an ultra-high-resolution scale (Sartorius Micro SC 2, Göttingen, Germany) with a precision of 0.0001mg. Calibration was carried out using a 2.001mg calibration weight ensuring the scale’s accuracy. To establish a consistent starting temperature, two 0.2mL plastic tubes were cooled on ice for five minutes. Each tube was weighed three times. Following this, an ant was placed inside each tube, and the tubes were again cooled on ice for five minutes to minimise ant movement. The tubes were then weighted three times to determine ant weight. After this, the ants were allowed to recover for five minutes before being placed in the volume estimation setup. Ants were provided either 0.5M or 1M sucrose solution *ad libitum*. Post-recording, the ants were returned to the same 0.2mL tubes and put on ice for another five minutes before undergoing their final weighing. At the end of each experimental day, all ants used were reintegrated into their respective colonies. In order to compare the weight of solution consumed by each individual with the estimated volume consumed, the density of the sucrose solutions used was determined. 1.5mL plastic tubes were placed on ice for five minutes, and later weighed three times. Using a precise repetitive pipette HandyStep® touch (Brand GmbH, Wertheim, Germany), the tubes were then filled with 0.5mL of the solution, cooled on ice for another five minutes, and subsequently weighed three times. Using the average density of each solution, the weight of solution consumed by each individual was then converted to volume. The density of the 0.5M sucrose solution was measured as 1.06 ± 0.02 g/mL (N = 17) and that of the 1M solution as 1.11 ± 0.04 g/mL (N = 22).

#### Experimental application - Volume estimation setup

Our setup (Figure 3) consisted of a resin 3D-printed platform (Ø 45mm, height 5mm) filled with water to prevent the ants from escaping. Within the platform, there was an internal structure (height 7mm) with a maximum width of 11mm at its broadest point. This structure tapered down to a 3mm width channel that extended a length of 6mm, ultimately leading to a well (Ø 4.5mm) where the solution being tested was placed. This entire setup was attached to two fixed Raspberry Pi HQ camera systems equipped with 6mm wide-angle lenses (Raspberry Pi Foundation, Cambridge, United Kingdom). One camera captured a top-view of the ant whilst the other recorded a side-view. The cameras were synchronized by implementing a client-server architecture within the same network, ensuring simultaneous triggering. Focal ants were placed on the largest part of the internal structure and recorded for the entirety of the drinking event having access to *ad libitum* solution. Once full, the ant was removed from the platform and returned to its respective colony. Using this setup, we provided the ants with both a range of sucrose solutions, as well as a gradient of caffeine-laced sucrose solution, in this way quantifying the ant’s feeding response to different food sources.

**Figure 3.**
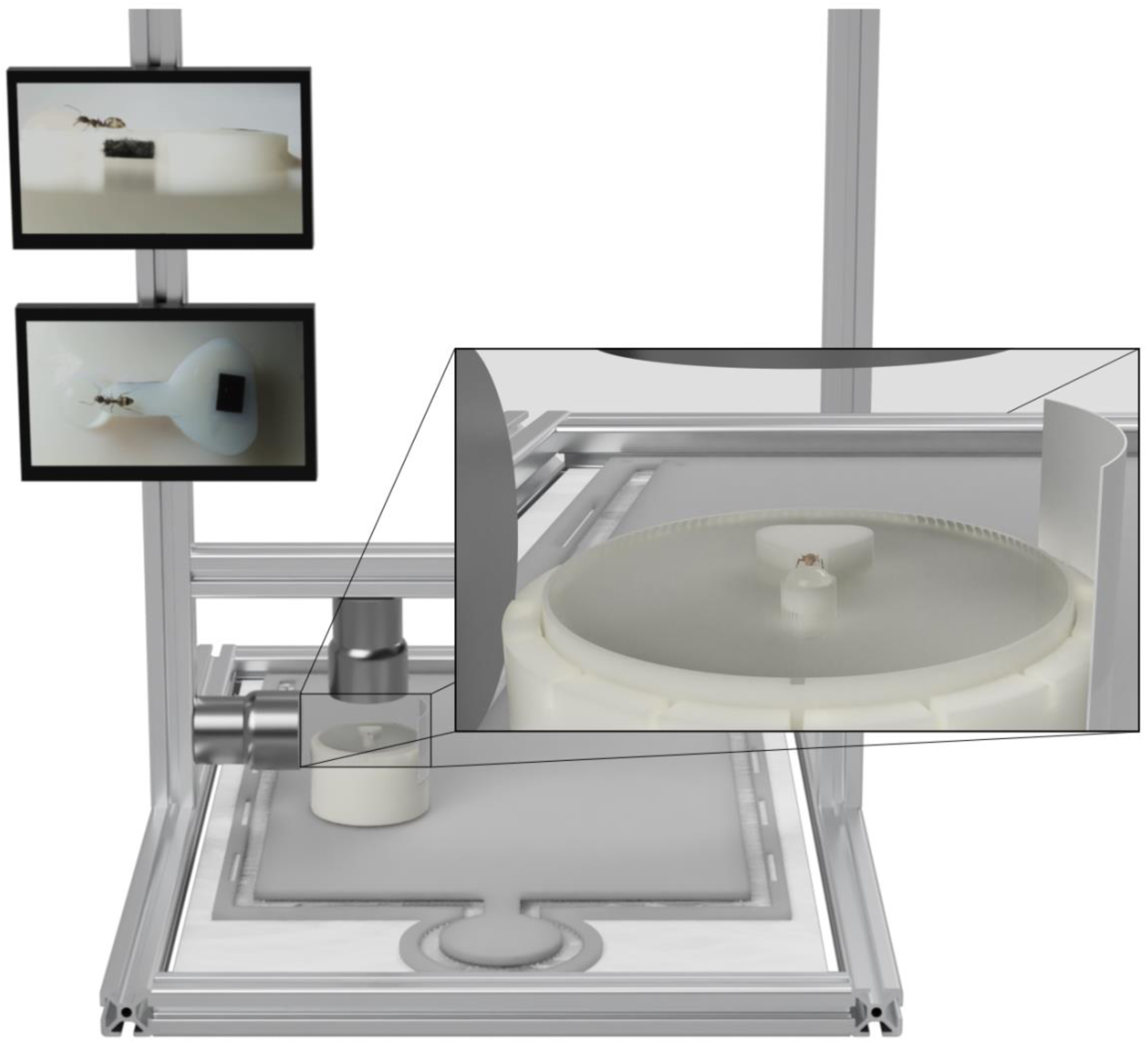
Volume estimation experimental setup. Resin 3D-printed platform (Ø 45mm, height 5mm) filled with water. Within the platform, an internal structure (height 7mm) with a maximum width of 11mm at its broadest point. This structure tapers down to a 3mm width channel that extends a length of 6mm, ultimately leading to a well (Ø 4.5mm) where the solution being tested is placed. The entire setup is attached to a fixed dual-camera system equipped with 6mm wide-angle lenses recording simultaneously a top-view and a side-view.

#### Data extraction and processing

In total 571 ants were recorded across all experiments. However, 146 recordings were discarded from the analysis (specified in the detailed statistical analysis and code), predominantly due to incorrect volume estimation resulting from the ant drinking at non-optimal positions. These ants were identified during the use of the GUI which allows the user to select initial and final time points for the feeding event. To exclude these, we simply did not select these time points, thus saving them automatically as empty values, making it easy to exclude them. As detailed above, all recordings were processed using DeepLabCut 3D pose estimation, and both crop load and consumption rate was obtained for each individual by fitting a linear regression to the estimated volumes over time. Overall, 114 ants were analysed for the accuracy measurements (experimental validation), and the seven volume estimation approaches proposed used. For the method’s experimental application, 311 ants were fed either pure sucrose or caffeine-laced sucrose dilutions, and their consumption was estimated by using the HVSpline method exclusively.

#### Statistical analysis

The complete statistical analysis output, and the entire dataset on which this analysis is based, is available from Zenodo (https://doi.org/10.5281/zenodo.11655204).

All plots and statistical analysis were generated using R version 4.2.2 (Wickham, 2016, 2022; R Core Team, 2022). All measurements were analysed using linear mixed-effects models (Bates *et al*., 2015). DHARMa (Hartig, 2022) was used to assess linear model assumptions and MuMIn (Bartoń, 2022) to obtain a measure of goodness of fit. Analysis of variance tables were used to test the effects of the regression’s coefficients (Fox & Weisberg, 2019). Estimated marginal means and contrasts were obtained using the emmeans (Lenth, 2022) package with Bonferroni adjusted values accounting for multiple testing. We avoid the use of p-values, and their associated binary decision of significant/nonsignificant, instead reporting effect size estimates and their respective 95% confidence intervals shown throughout the results section as (estimate [lower limit, upper limit, N = sample size]).

## Results and Discussion

### Experimental validation – Accuracy measurements

As stated in the introduction, traditional methods of quantifying consumption are subject to considerable error at small volumetric scales. This makes it extremely difficult to establish a ground truth value and thus to estimate the accuracy of new methods. Nevertheless, here we attempt to do so by assessing the agreement between gravimetric measurements, and each of the seven proposed methods of estimating ingested volume (Figure 4). The Ellipsoid volume estimation method agreed most with weight, albeit producing consistently lower estimates, on average, by 62nL [45nL, 79nL, N = 728]. In part, this is due to the scale being particularly prone to error at small values, and due to the necessary density conversion. Moreover, weight measures the mass an individual gained or loss, regardless of if this was due to ingestion or simply due to a part of the solution being attached to the individual’s body. At larger scales, this is often negligeable, but at the nanolitre scale a small drop of solution on the individual’s body will lead to significant deviations from accurate measurements. This is highlighted by the strong positive correlation between the difference of weight and volume estimation and their mean. Such correlation is driven by weight values being, for the most part, higher than estimated volumes (Figure 4).

**Figure 4.**
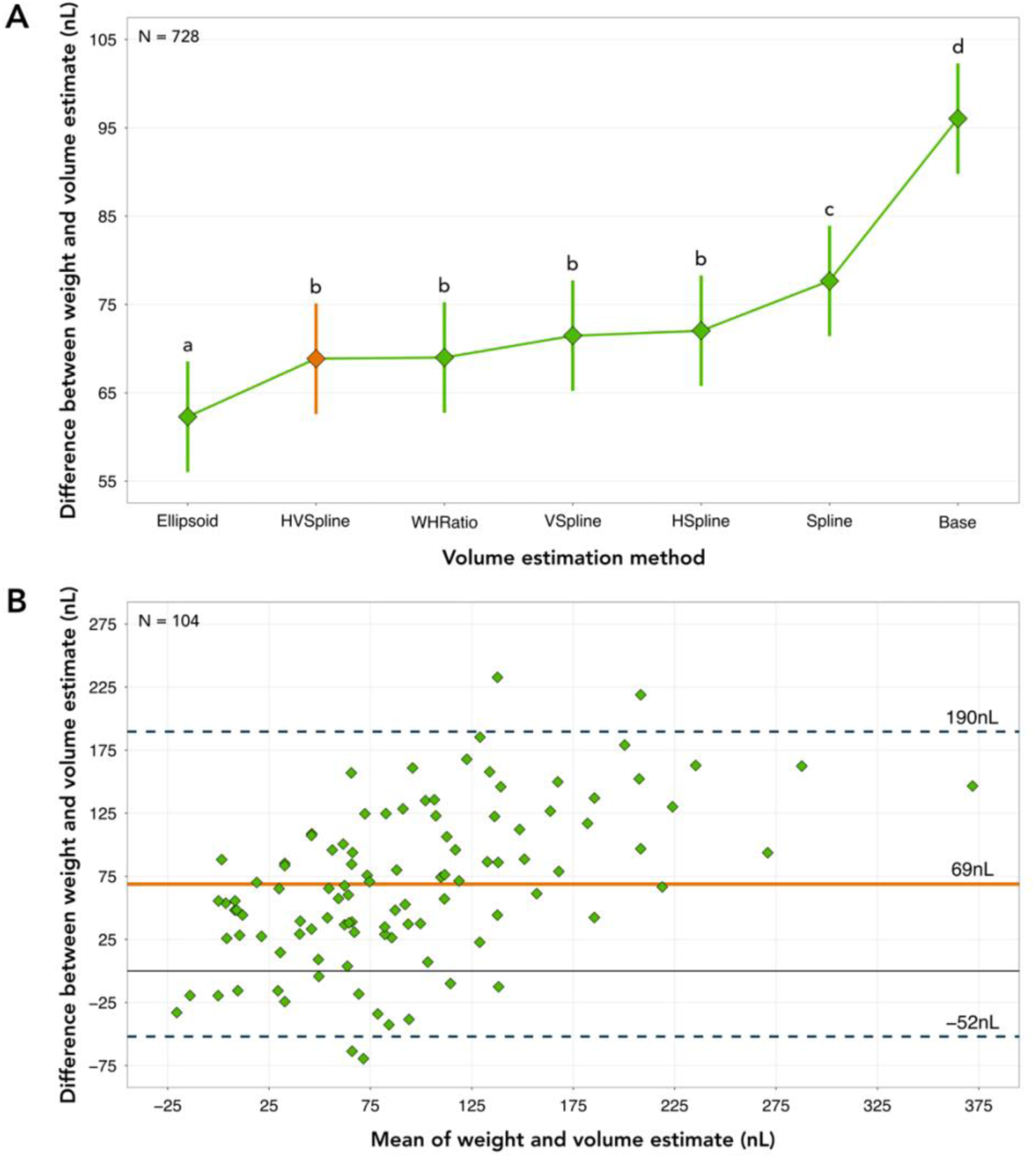
Comparison between the volume estimates obtained from each of the methods proposed, against a gravimetric measure of consumption. **(A)** Accuracy of volume estimates compared to weight measurements. Diamonds represent the estimated marginal means obtained from the linear mixed-effects model and whiskers the respective standard errors. Letters reflect statistical differences between the agreement of each volume estimation method and the weight measurement based on the estimated confidence intervals. The Ellipsoid method agrees best with weight (62nL [45nL, 79nL], N = 728). However, we optted for the HVSpline method (69nL [52nL, 86nL], N = 728), as weight measures have significant associated error, and this volume estimation method can be used with a wider variety of shapes. **(B)** Bland–Altman plot for the agreement between weight measurements and the HVSpline volume estimates. Diamonds represent individual feeding events, and the orange line the average difference between the two methods, used as an estimate of accuracy. Dashed lines represent the 95% limits of agreement. Note that there is a strong positive correlation between the mean of the two measurements and their difference, which is predominantly driven by the weight measurements (see Statistical Analysis and Code). Moreover, in almost all cases, the weight measurement is higher than the estimated volume.

In addition, we compared the estimated volumes with a solution of known volume, 138nL, which was consumed in its entirety (see Statistical Analysis and Code). In this case, the Ellipsoid volume estimation method also agreed best with the known volume, albeit producing on average 53nL [24nL, 81nL, N = 70] lower estimates. Importantly, at the nanolitre scale, a liquid drop suffers from large evaporation rates. Thus, the discrepancy between the estimated volume and the available solution is likely due to evaporation. The exact amount of solution evaporated cannot be quantified as we did not control for the time it takes for ants to first contact the drop. However, we estimate that ants took between two and five minutes to reach the solution and therefore expect an absolute minimum evaporation loss of 25nL (see Statistical Analysis and Code).

Even though the Ellipsoid method best agrees with both weight and a solution of known amount, we opted for the second highest agreeing method, the HVSpline. This is because, whilst an ant gaster, due to its natural shape, is correctly approximated to an ellipsoid, such approximation is not reasonable for most other organisms. Furthermore, since weight measurements are likely to be overestimating consumption, opting for the most agreeing volume estimation method might not be accurate, as in principle, such method would likely also overestimate consumption. The HVSpline method maintained a comparably high level of agreement to weight (69nL [52nL, 86nL, N = 728]) and known solution volume (55nL [27nL, 84nL, N = 70]) in comparison to the Ellipsoid method, whilst relying on fewer *a priori* assumptions of the organism’s shape, being therefore more transferable to other organisms.

### Experimental application – Higher sucrose concentrations increase crop load but decrease the consumption rate of an invasive ant

To assess if the method proposed is able to detect known behavioural patterns, we applied it to Argentine ants feeding on a wide range of sucrose solutions. As sucrose concentration increases, so does the energy provided by the solution. However, with an increase in sucrose, the viscosity of the solution also increases. Ants ingest food through a sucking-pump system which creates negative pressure through muscle contraction, driving fluid into the mouth (Paul, Roces & Hölldobler, 2002; Falibene, Rössler & Josens, 2012). In this way, ants are able to vary their consumption rate (Josens & Roces, 2000), for example by increasing pumping frequency, whilst maintaining a constant volume taken per pump contraction (Falibene & Josens, 2008; Falibene, Gontijo & Josens, 2009). Higher viscosity makes it harder for ants to ingest food, leading to longer feeding times to reach the same crop load. Interestingly, if viscosity is artificially manipulated, and a low sucrose concentration solution is made to be highly viscous, ants rapidly decrease their pump frequency, resulting in demotivation for the food source (Lois-Milevicich, Schilman & Josens, 2021). In some species, high viscosity solutions can even trigger a switch from trophallaxis – where ants ingest a liquid and later share it with their nestmates – to pseudotrophallaxis – where ants hold a liquid drop between their mandibles through surface tension and later share it with their nestmates without ingestion (Fujioka, Marchand & LeBoeuf, 2023).

As expected, our data suggests that an increase in sucrose concentration results in a relative increase of crop load, further reinforcing that individual ants are capable of regulating their food intake (Figure 5). Ants collecting diluted solutions reach lower crop loads (0.125M: 306nL [177nL, 435nL, N = 269]; 0.25M: 278nL [151nL, 405nL, N = 269]) than those provided higher molarity solutions (0.5M: 360nL [235nL, 486nL, N = 269]; 1M: 361nL [235nL, 487nL, N = 269]; 2M: 379nL [254nL, 504nL, N = 269]). A similar pattern has been previously observed for Argentine ants, where crop load increases with sucrose concentration, albeit with such increases being lost at concentrations higher than 2M (Sola & Josens, 2016). Moreover, crop load volumes obtained in our experiment are generally higher than those previously reported, which reach a maximum around 200nL (Sola, Falibene & Josens, 2013; Sola & Josens, 2016). Such differences could be due to previous studies using Argentine ants from their native range, whilst we used ants from their invasive range, which could potentially influence feeding behaviour. Additionally, the ants used in our study were kept in considerably smaller colonies and starved for a shorter period of time when compared to previous work. Feeding patterns are highly variable and dependent on multiple factors. For example, in the ant *Camponotus mus*, increases in sucrose concentration were shown to increase crop load (Josens, Farina & Roces, 1998), with feeding varying with colony starvation (Josens & Roces, 2000).

**Figure 5.**
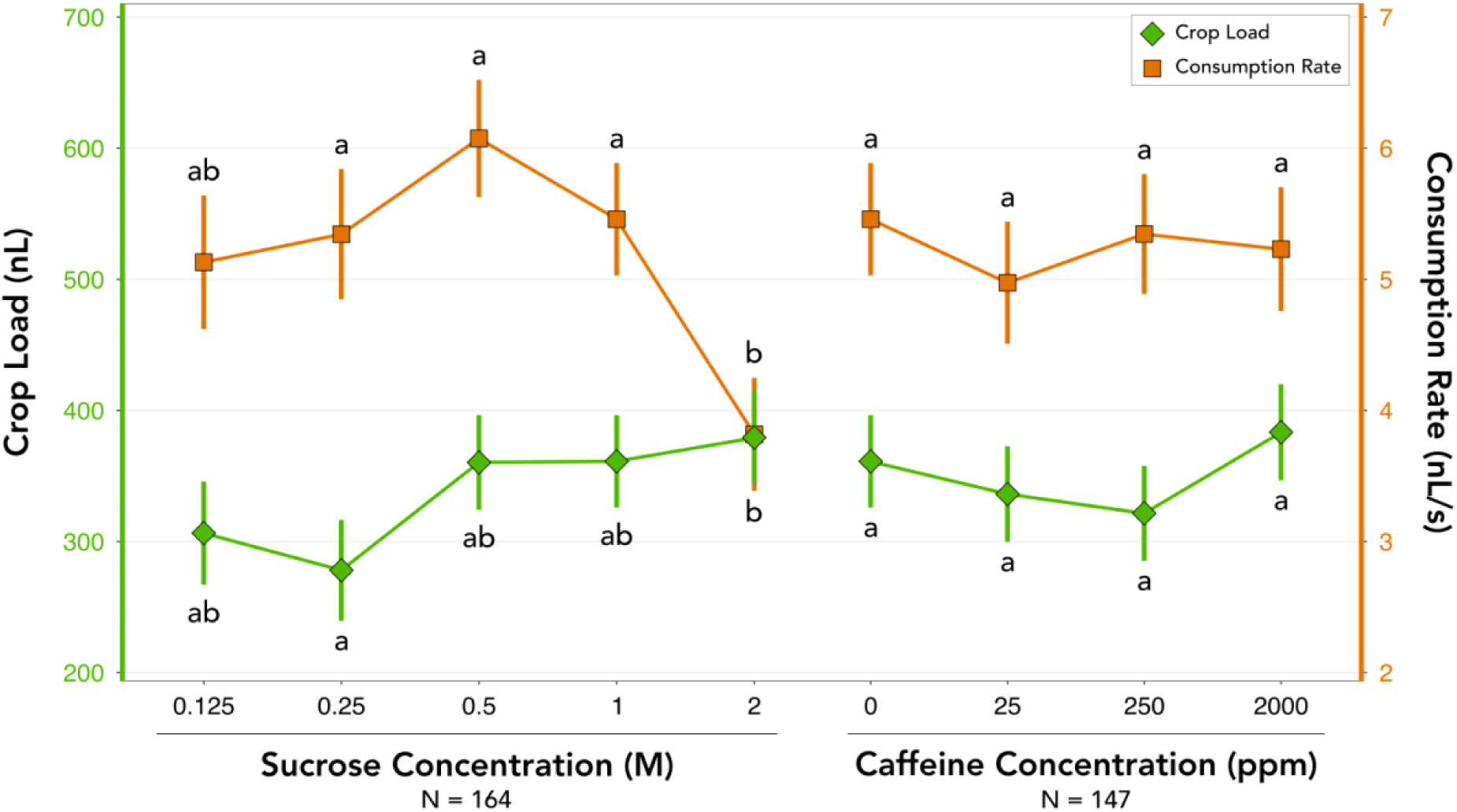
Quantifying invasive ant feeding behaviour by three-dimensionally reconstructing their body during feeding. Diamonds and squares represent the estimated marginal means obtained from the linear mixed-effects models and whiskers the respective standard errors. Letters reflect statistical differences between treatments which are connected based on the estimated confidence intervals. Similarly to what can be found in literature, we find a trend towards increased crop load with increasing sucrose concentrations, with consumption rate decreasing as sucrose concentration increases. This reflects the trade-off between solutions which have high sucrose molarity having high energy output whilst being hard to drink from due to high viscosity. Overall, the presence of caffeine appears to have no effect on either crop load or consumption rate. This suggests ants are not averse to it or simply cannot detect its presence.

Consumption rate (Figure 5) was relatively constant at lower sucrose concentrations (0.125M: 5.1nL/s [3.6nL/s, 6.6nL/s, N = 269]; 0.25M: 5.3nL/s [3.9nL/s, 6.8nL/s, N = 269]), reaching its maximum at 0.5M (6.1nL/s [4.7nL/s, 7.4nL/s, N = 269]) and decreasing at high concentrations (1M: 5.5nL/s [4.1nL/s, 6.8nL/s, N = 269]; 2M: 3.8nL/s [2.5nL/s, 5.2nL/s, N = 269]). Importantly, the consumption rates obtained in our experiment are almost five-fold those reported in Sola and Josens (2016) for the same species, which reach a maximum around 1.6nL/s. However, in this previous work, feeding time was taken as the time during which the ant’s mandibles touched the solution, an indirect measure of ingestion, and these were considerably longer than those we observed. Interestingly, between experiments ants were kept on 0.5M sucrose solutions which could explain the peak consumption rate at this concentration. This is expected, as ants are more likely to be demotivated by solutions which are more diluted than those they have previously experienced (Cassill & Tschinkel, 1999; Wendt *et al*., 2019), whilst being physically slowed down by the high viscosity of higher concentration solutions (Lois-Milevicich, Schilman & Josens, 2021).

### Experimental application – Caffeine presence has no effect on invasive ant feeding behaviour

Additionally, we took the opportunity to feed the ants a range of caffeine-laced sucrose solutions. Caffeine has been reported to decrease Argentine ant foraging times, likely due to its beneficial cognitive effects. For this reason, caffeine was suggested as a suitable additive to slow-acting baits, as it might lead to increases in recruitment and consumption, thus potentially improving current invasive ant management (Galante *et al*., 2024). Understanding the reason for this improvement is important, as for example, caffeine could simply be improving food palatability. Thus, it is important to study if caffeine has an effect on the feeding behaviour, either the total volume ingested or the rate at which it is ingested, in order to assess if the reported neuroactive effects of caffeine are in part a result of increased preference and motivation for caffeine-laced solutions.

The presence of caffeine (Figure 5) had no clear effect on either crop load (0ppm: 361nL [235nL, 487nL, N = 269]; 25ppm: 336nL [208nL, 464nL, N = 269]; 250ppm: 321nL [195nL, 448nL, N = 269]; 2000ppm: 383nL [257nL, 510nL, N = 269]) or consumption rate (0ppm: 5.5nL/s [4.1nL/s, 6.8nL/s, N = 269]; 25ppm: 5.0nL/s [3.5nL/s, 6.4nL/s, N = 269]; 250ppm: 5.3nL/s [3.9nL/s, 6.8nL/s, N = 269]; 2000ppm: 5.2nL/s [3.8nL/s, 6.7nL/s, N = 269]). This suggests caffeine has no effect on reward value perception, either because ants can’t detect its presence or are indifferent to it. Thus, the addition of caffeine to toxic baits is likely to result in foraging improvements driven by the alkaloid’s effect on learning and memory rather than on motivation. On average, individual ants weighed 0.5mg ± 0.1mg (N = 145), meaning ants under the 25ppm of caffeine treatment ingested on average 16.8mg of caffeine per kilogramme of body mass [10.4mg/kg, 23.2mg/kg] and those under the 250ppm treatment 160.5mg/kg [97.5mg/kg, 224mg/kg]. For reference, humans ingest under 10mg/kg per day at most (Verster & Koenig, 2018).

### System strengths, limitations, and potential improvements

The method proposed allows researchers to quantify the feeding behaviour of small invertebrates, not only when it comes to the total amount of food ingested, but also the rate at which food is ingested. It takes advantage of the fact that the bodies of some animals expand while feeding, and three-dimensionally reconstructs their body over time, and thus reducing the number of approximations required to estimate ingested volume.

The method is high throughput, with recordings taking on average 3.51 ± 0.87 minutes (N = 571) per individual and ants drinking on average 1.5 ± 1.1 minutes (N = 466), which is similar to previously reported average drinking times of 1.4 minutes for the same species (Galante *et al*., 2024). The pose estimation process can be time consuming, depending on the available computational resources, but, for the most part, it does not require human intervention. We estimate that data processing, using the GUI, allowed us to look at over 40 videos per hour. Moreover, the system is affordable, as it can be assembled with different materials, such as cameras which are already available, or built with Raspberry Pi HQ camera systems which are extremely affordable and convenient.

One limitation of our specific system was that the accuracy of the tracked points, and thus of the three-dimensional reconstruction and volumetric measurements was compromised when the individual ant wasn’t correctly positioned in reference to the cameras. This led us to exclude a large number of recordings for which quantifying feeding behaviour was not possible. In our case, this was mitigated by a high sample size. However, if possible, building a recording system with more cameras would mitigate the need for the animals to be in specific positions, enabling for example the tracking of animals mid-flight (Maya *et al*., 2023; Håkansson *et al*., 2024). This could overcome limitations such as occlusions, perspective distortions or ambiguity in depth estimation. Importantly, whilst the system is internally consistent, meaning the results of different treatments are always comparable to each other, external consistency, and thus comparison across different experiments, will rely entirely on the standardisation of the reference points and the accuracy of the camera calibration. To overcome this, we suggest, for example, using camera lenses with microscope calibration scales.

During our accuracy measurements for the experimental validation, we were able to directly compare our method with the traditional gravimetric approach. We found that weighing was not only more time consuming, especially as usually three replicates for each weighing are required, but also resulted in a substantial number of individuals being lost due to excessive anaesthesia or during transfer to the scale. Interestingly, ants which were anesthetised and used in the gravimetric accuracy measurements, had a lower crop load (max = 299nL, N = 104) and consumption rate (max = 4.1nL/s, N = 104) than those estimated for similar sucrose solutions (Figure 5) which were not anesthetised and weighed (0.5M: 360nL at 6.1nL/s; 1M: 361nL at 5.5nL/s). This is likely due to the stress induced by cooling anaesthesia further reinforcing the benefits of a non-invasive method which does not require anaesthesia. In fact, ants which were anesthetised, either by cooling or carbon dioxide, were previously shown to be less willing to feed on sugary solutions (Mailleux, Deneubourg & Detrain, 2000).

The measuring device proposed here is extremely useful for the quantification of feeding behaviour in ants, especially invasive ones, which tend to be small yet expand during feeding. For Argentine ants, consumption rate was for the most part linear. However, the method is flexibly applicable to animals whose feeding dynamics are not constant over time, by simply fitting a different model to the data. We predict the system would also be useful for disease vector studies, for example, elucidating feeding patterns in mosquitoes, potentially leading to new control methodology. Importantly, the fact that the method is non-invasive, means it is also suitable for larger insects, which could be reliably weighed, but in this way would be less disturbed, thus making it easier to capture their natural behaviour, even in field settings if required. Finally, in principle, this system could also be applied to non-living objects, for example allowing the measure of volumetric changes in medical applications, where equipment usually has a high cost.

Overall, the proposed method of directly estimating volumetric changes over time is much faster and less invasive than most currently used methods. Considering it is currently almost impossible to establish a ground truth measurement of the feeding behaviour of small invertebrates, we believe we successfully demonstrated the method is both accurate and capable of detecting known behavioural patterns. Additionally, we demonstrate its potential to measure feeding preference and chemical perception. Ultimately, this could provide important insight on the preferences of disease vectors, such as mosquitoes feeding on different blood types, and have direct impacts on invasive ant management strategies.

## Acknowledgements

We thank E. Sequeira and S. Abril for ant collection, L. Aichner and L. Guyton for help with data collection, and S. Ibarra for ant illustrations.

## Ethical statement

We have conducted all experiments in accordance with the guidelines that are applicable to working with the model organism in the European Union. Colonies were kept in closed boxes under oil baths in order to prevent any escape.

## Author contributions

**H. Galante:** Conceptualization, Methodology, Software, Validation, Formal analysis, Investigation, Data Curation, Writing - Original Draft, Visualization, Supervision. **T. J. Czaczkes:** Resources, Writing - Review & Editing, Supervision, Project administration, Funding acquisition. **M. De Agrò:** Conceptualization, Methodology, Validation, Writing - Review & Editing.

